# Prior College Biology Coursework Increases Student Outcomes for Underrepresented Minority Students studying Anatomy and Physiology

**DOI:** 10.1101/2023.02.23.529571

**Authors:** Brian R. Stephens

## Abstract

Student performance and achievement in anatomy and physiology (AP) are determining factors in admissions into medical and other health-allied programs. Madison College has had historically poor course retention and pass rates for underrepresented minorities in A&P. Course and enrollment data from 2013 – 2020 were examined to determine whether students who optionally took lab-based biology coursework had improved student outcomes. Underrepresented minority students who completed ≥ 1 lab-based college biology course before entering AP had improved student outcomes evidenced by an increased pass rate and a reduced failure rate. Statistical analysis uncovered a statistically significant association (*p*≤ 0.10) with prior lab-based college biology coursework and student outcomes in AP for underrepresented minority students.

## INTRODUCTION

Anatomy and physiology is a challenging gateway course characterized by high failure and attrition rates (Hopp, 2009; Russell, Young, & Lehning, 2016; Slominski, Grindberg, & Momsen, 2019). Russell et al. reported in a large, multi-year (2003 – 2015) study a combined failure and withdrawal rate of 41% for AP students (Russell, Young, & Lehning, 2016). Similar studies indicate a combined failure and withdrawal rate ranging from 20% to 50% (Hopp, 2009; Russell, Young, & Lehning, 2016; Rosenzweig, 2012). Why is AP so difficult? It’s not entirely clear. Sturges and Maurer (2013) reported both faculty and students felt the discipline itself (e.g., memorizing numerous terms, understanding how biological systems together, additive nature of anatomy and physiology) was the primary driver for perceived course difficulty. In another study, Slominski et al. (2019) found similar results and found that anatomy and physiology is difficult not due to deficiencies in teaching, instruction, or student learning approaches but due to the inherent difficulty of the discipline itself.

Due to the difficulty of the course, the ideal student entering AP should have a solid framework of basic biology from high school. This means that students should possess knowledge of the cell cycle, the location and function of organelles, Mendelian inheritance, the location, appearance, function of body tissues, a basic understanding of organ systems, enzymes, and cellular respiration (Khalil, Lazarowitz, & Hertz-Lazarowitz, 2014). Students should be able to handle a microscope and have familiarity with dissections and lab safety; students should also be familiar with the self-directed nature of biology labs and biology practicals (Khalil, Lazarowitz, & Hertz-Lazarowitz, 2014). Given the well-known disparities in education in the United States (Smedley, Stith, Colburn, & Evans, 2001), biology prerequisites could potentially provide valuable knowledge and lab skills to students who either may need a refresher or never had an opportunity in high school.

The “achievement gap” typically refers to the disparity in achievement between white and underrepresented minority students. For example, Wisconsin has highest achievement gap in the nation for African-American and white students in math and reading for 4^th^ and 8^th^ graders as well as high school graduation rates (NAEP, 2020). Over the last decade, the education system in the State of Wisconsin has consistently ranked last or near last on most measures of equity between white and underrepresented minority students (Becker, 2015; NAEP, 2020; Kremer, 2021). While these findings can potentially motivate well-intentioned reform, Ladson-Billings (2006) believes the dialogue around “achievement gaps” is controversial and is blunt in her assessment of it. She believes the discourse of achievement gaps places academically struggling students in a vacuum without consideration of factors important to student success such as disparities in health, wealth, and school funding (Ladson-Billings, 2006; 2007). Ladson-Billings (2006) posits that the observed “gap” between white and underrepresented minority students isn’t due to a lack of achievement, rather, due to an accumulated “education debt” owed to underrepresented minorities. The author believes this educational debt has historical, economic, sociopolitical, and moral underpinnings. For example, Ladson-Billings (2006) points to numerous historical examples of excluding underrepresented minorities from education including the Lemon Grove Incident, *Plessy v. Ferguson*, and *Mendez v. Westminster*; further, Ladson-Billings (2006) points out that African-American students in the South did not have universal secondary schooling until 1968 (Wright, 1941; Love, 2004); and that *No Child Left Behind* left a mandate to annually test students but failed to invest adequate resources in public education (Imazeki & Reschovsky, 2004). After painting this picture, Ladson-Billings (2006) rhetorically asks: “Why, then, would we not expect there to be an achievement gap?”

Milner (2012) also critiques the national discourse on “achievement gaps” on following grounds. First, like Ladson-Billings (2006) he believes explanations on achievement gaps focus on differences between underrepresented minority students and white students without examining reasons behind those disparities (Milner, 2012; 2013). Second, such comparisons tend to frame white students as the standard to which all other students are compared (Milner, 2012). For example, Asian students have been described as “outperforming” in STEM, here, it is implicit that Asian students are being compared to the norm: white students (Milner, 2012; Chen & Buell, 2018). Third, discussions on “achievement gaps” can frame underrepresented minority students as culturally and intellectually inferior (Milner, 2012). Lastly, these comparisons tend to focus mostly, if not entirely, on the individual or the ethnic group(s) as opposed to the systemic inequality, racism, and sexism that lead to these achievement gaps (Milner, 2012).

What are some well-known disparities in education? The Office of Civil Rights within the U.S Department of Education collects data on all public schools (pre-K – 12^th^ grade) in the United States including D.C. and U.S territories. This includes charter schools, alternative schools, juvenile justice, and special education facilities. The Office of Civil Rights reports that schools that are predominately African-American, Hispanic, and Native American students have limited access to high-level math and AP science courses and experience increased rates of being held back (or retained a year) (CRDC, 2014; Klopfenstein, 2004). Further, the Office of Civil Rights reported that African-American students were expelled, suspended, referred to law enforcement, and transferred to alternative schools at more than twice their share of total enrollment (CRDC, 2021); indeed, between 2011 – 2012 the State of Wisconsin led the nation in the suspensions of African-American students (Losen, Hodson, Keith II, Morrison, & Belway, 2015) Unfortunately, this disparity in treatment is entrenched before the children learn to read. For example, between 2017 – 2018, African-American boys were a meager 9.6% of total enrolled pre-K students (2 ½ to 4 ½ years old) in the United States yet made up 34.2% and 30.4% of all pre-K suspensions and expulsions, respectively (CRDC, 2021)

In Fall 2020, Madison College launched an <u>Equity and Inclusion plan,</u> which, among other things, tasked departments to examine courses and prerequisites; the plan also encouraged analysis of shortcomings in closing equity gaps as well as developing strategies to attract, train, and retain underrepresented minority students in STEM and allied-health fields. Because success in AP is important in achieving these goals, two questions were asked. First, how well do underrepresented minority students succeed in AP? Two, to what extent does the literature inform educators, biology departments, and administrators of strategies to improve to the success of underrepresented minority students in AP?

There is a paucity of research examining these two questions. Indeed, a review of the literature yielded one article published in this journal. In 2007, Beeber and Biermann reported that the AP pass rate for “at-risk” students substantially increased after enrolling and passing a college biology course. What was particularly striking from the findings is that a general biology course was superior to a specially designed “biology foundations” course created to help students succeed in AP. This was interesting because it suggests two things. First, reinventing the wheel and creating specialized classes may not be necessary to improve student outcomes for underrepresented minorities in AP; second, other lab-based college biology courses may provide the preparatory background, confidence, and knowledge to increase student success in AP. While the conclusions of Beeber and Biermann (2007) provide a framework for further study, three gaps persist that merit pause and consideration. First, Beeber and Biermann (2007) referred to students at a majority-minority community college as “at-risk” which can be a vague (if not loaded) term. Second, Beeber and Biermann (2007) did not provide precise demographic or educational characteristics of their sample or statistical analysis for their data. Third, Beeber and Biermann (2007) considered students who withdrew from the course in their analysis. There are valid reasons a student may withdraw from AP independent of course content or course difficulty; for example, a student may withdraw due to work, personal, or family obligations (Hall & Smith, 2003). To that end, this study examines students who withdrew as well as those who persisted and incorporates some statistical analysis of the data to help better inform educators, biology departments, and administrators.

## METHODS AND STATISTICAL ANALYSIS

Established in 1912, Madison Area Technical College (or Madison College) is a two-year community college located in Madison, Wisconsin. Madison College offers two introductory AP courses with historically poor course retention and success rates for underrepresented minority students. The first introductory course is General Anatomy and Physiology, which is required for admission into occupational assistant, medical and surgical laboratory technician, radiography, respiratory therapy, and dental hygienist programs. The second introductory course is anatomy and physiology 1 (or AP1), which is the first of a two-semester sequence and is required for admission into nursing and medical programs. I sought to determine whether underrepresented minorities who optionally took a college biology course with a lab component had improved course outcomes in AP. I also wanted to determine whether there was a statistically significant association between pass and fail rates for students who completed AP <u>and</u> did not withdraw from the course.

From Summer 2013 – Fall 2020, approximately 6,087 students enrolled two introductory courses at Madison College. Out of these students, only the 921 whom self-identified as underrepresented minority students were included in this study. For the purposes of this study, underrepresented minorities were defined as those as identify as African-American, Hispanic, and Native American. From this group, data was collected on prior lab-based college biology coursework, final grades, demographics, age, sex, external GPA, and standardized test scores. External GPA and standardized test scores were not available for all students.

Lab-based college biology courses included Introduction to Human Biology, Introduction to Zoology, and Botany. Unlike most institutions, Madison College does not have a “C-”, thus, passing was defined as a student receiving a “C” or better while any “D” or “F” was considered failing, and any type of “W” was considered a withdrawal. Students taking the course for credit or pass fail were omitted from the study.

Data were evaluated using Excel (Office 365 version 2008) and SPSS (SPSS Statistics 27.0). Chi-square test of independence was used to make comparisons between groups using SPSS. Due to the preliminary nature of the study, the alpha level was set at .10. Therefore, p ≤ 0.10 is statistically significant.

## RESULTS

As indicated, **Table 1** shows the age and sex distributions of students enrolled in Anatomy and Physiology. The reported sex distribution was as follows: 100 males (28%) and 262 females (72%) for African-Americans, 4,133 females (80%) and 1,033 males (20%) for non-Hispanic Whites, 118 males (22%) and 417 females (78%) for Hispanic, and for Native Americans two (2) males (8%) and 22 females (92%). Values reported as mean ± SD. The average age of African-Americans was 27.8 ± 7.4 years (median: 27 years), non-Hispanic Whites were 24.9 ± 7.3 (median: 22.4 years), Hispanic was 23.4 ± 6.4 (median: 21 years), Native Americans was 25.3 ± 5.7 (median: 25 years). Students that didn’t disclose sex were omitted.

**Table 1:** Age and Sex Distribution of Students.

| | *n* | Mean Age ± SD | Median Age | Females (%) | Males (%) |
| --- | --- | --- | --- | --- | --- |
| **African-American** | 362 | 27.8 ± 7.4 | 27 | 262 (72%) | 100 (28%) |
| **Hispanic** | 535 | 23.4 ± 6.4 | 21 | 417 (78%) | 118 (22%) |
| **Native American** | 24 | 25.3 ± 5.7 | 25 | 22 (92%) | 2 (8%) |
| **Non-Hispanic Whites** | 5166 | 24.9 ± 7.3 | 22 | 4133 (80%) | 1033 (20%) |

As indicated, **Table 2** shows the reported external GPA and standardized test scores of students enrolled in the course. Because GPA and ACT test scores were not available for all students, only the students that had scores were calculated. For African-American students, the average external GPA (n=131) was 2.5 ± 0.6, the average Math ACT score (n = 43) was 16.3 ± 4.6, and the average English ACT score (n = 43) was 17.5 ± 2.6. For non-Hispanic white students, the average external GPA (n = 2,421) was 3.0 ± 0.6, the average Math ACT score (n = 1,539) score was 20.8 ± 4.2, and the average English ACT (n = 1,539) score was 20.7 ± 4.1. For Hispanic students, the average external GPA was 2.8 ± 0.6, the average Math ACT score (n = 181) score was 18.7 ± 3.4, and the average English ACT (n = 181) score was 17.8 ± 4.1. For Native American students, there was one external GPA score (n = 1) of 2.8 and no standardized test scores.

**Table 2:**
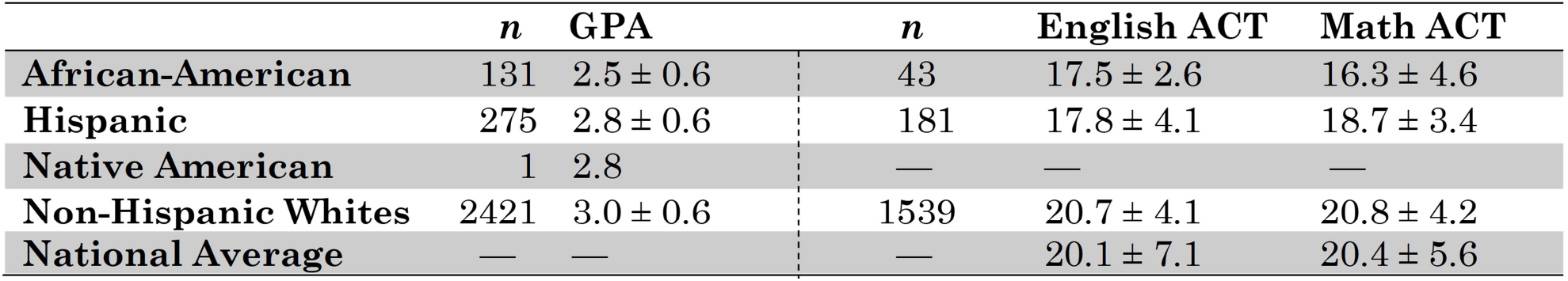
External GPA and Standardized Test Scores.

Next, students who had prior lab-based college biology coursework were compared with those who didn’t. Out of 6,087 students, 5,704 (94%) did not take a biology course before enrolling in anatomy and physiology, while only 383 students (6%) students did. Specifically, 6% (315/5,166) of non-Hispanic White students and 7% (68/921) of underrepresented minority students took ≥ 1 lab-based college biology course. The pass rate for underrepresented minorities increased 67.1% to 77.9% after completion of ≥ 1 college biology course. In addition, underrepresented minorities who took a college course decreased the failure rate from 14.5% to 4.4%, while non-Hispanic White student failures decreased from 7.6% to 5.4%. In contrast, pass rates of non-Hispanic White students appear to plateau from 82.3% to 82.2%. While the withdrawal rate decreased from 18.4% to 17.6% for underrepresented minorities, there was a nominal increase in the withdrawal rate from 10.1% to 12.4% for non-Hispanic White students (**Fig. 1**). Chi-square test of independence was used to determine whether there was a significant difference between the observed counts and expected counts. For the particular dataset in Fig. 1, while there was not a significant association between taking college biology and improved course outcome in AP for non-Hispanic White students (χ2 = 3.497, d.f. = 2, *p* = 0.17) (**Supplementary Fig. S1**), there was statistical significant association (p ≤ 0.10) for underrepresented minorities (χ2 = 5.801, d.f. = 2, *p* = 0.058) (**Supplementary Fig. S2**).

**Figure 1.**
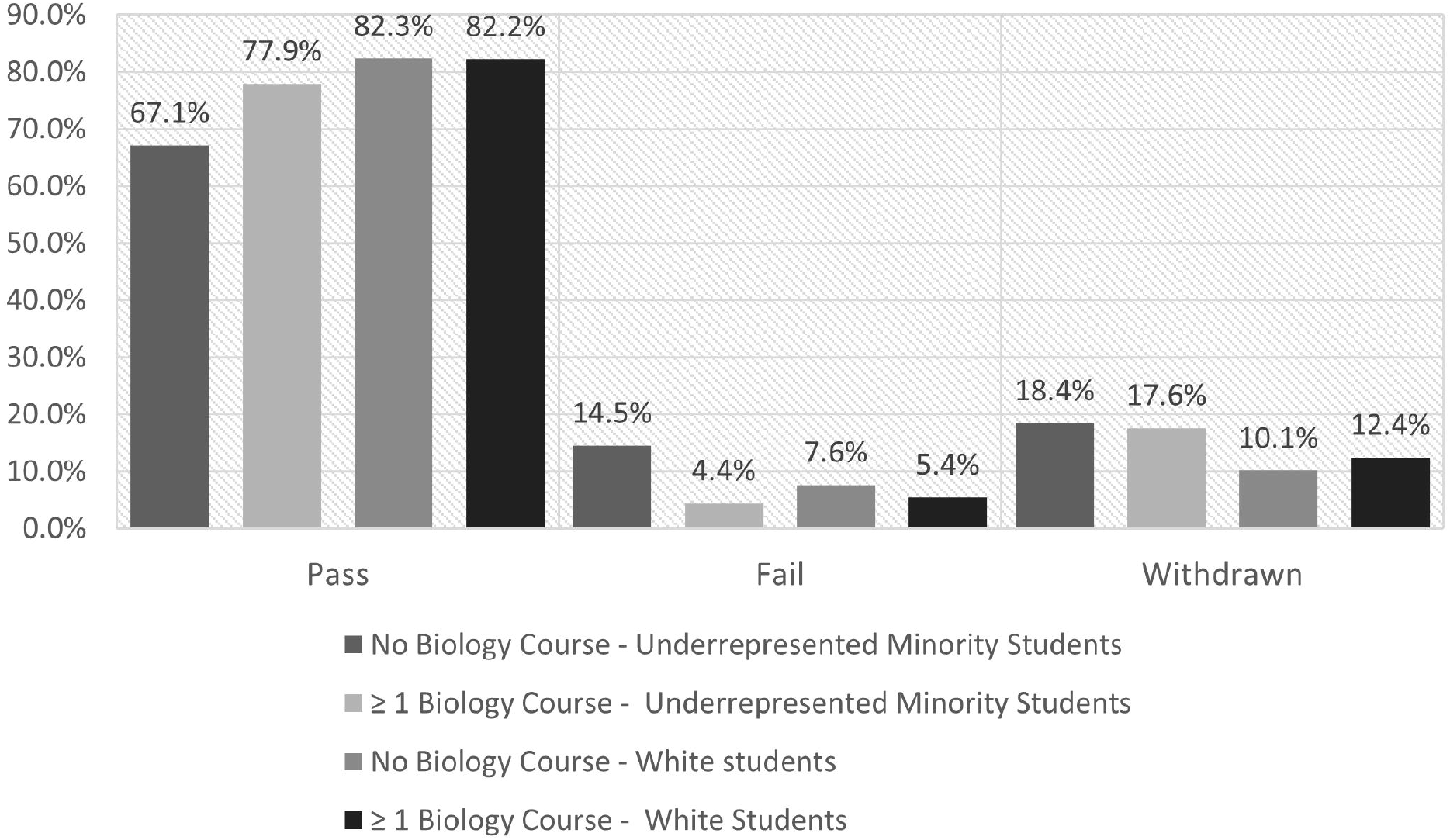

Since there can be compelling, non-academic reasons for a student to withdraw from a course, I next wanted to determine the pass and failure rates of students who completed AP <u>and</u> did not withdraw from the course. Of this group, underrepresented minorities taking ≥ 1 college biology course had an increase and decrease in the pass and failure rate, respectively (**Fig. 2**). Chi-square test of independence was used to determine whether there was a significant difference between the observed counts and expected counts. While there was not significant association between taking college biology and improved course outcomes AP for non-Hispanic White students (χ2 = 1.866, d.f. = 1, p = 0.10) (**Supplementary Fig. S3**), there was a statistically significant association for underrepresented minorities (χ2 = 5.732, d.f. = 1, *p* ≤ 0.01) (**Supplementary Fig. S4**).

**Figure 2.**
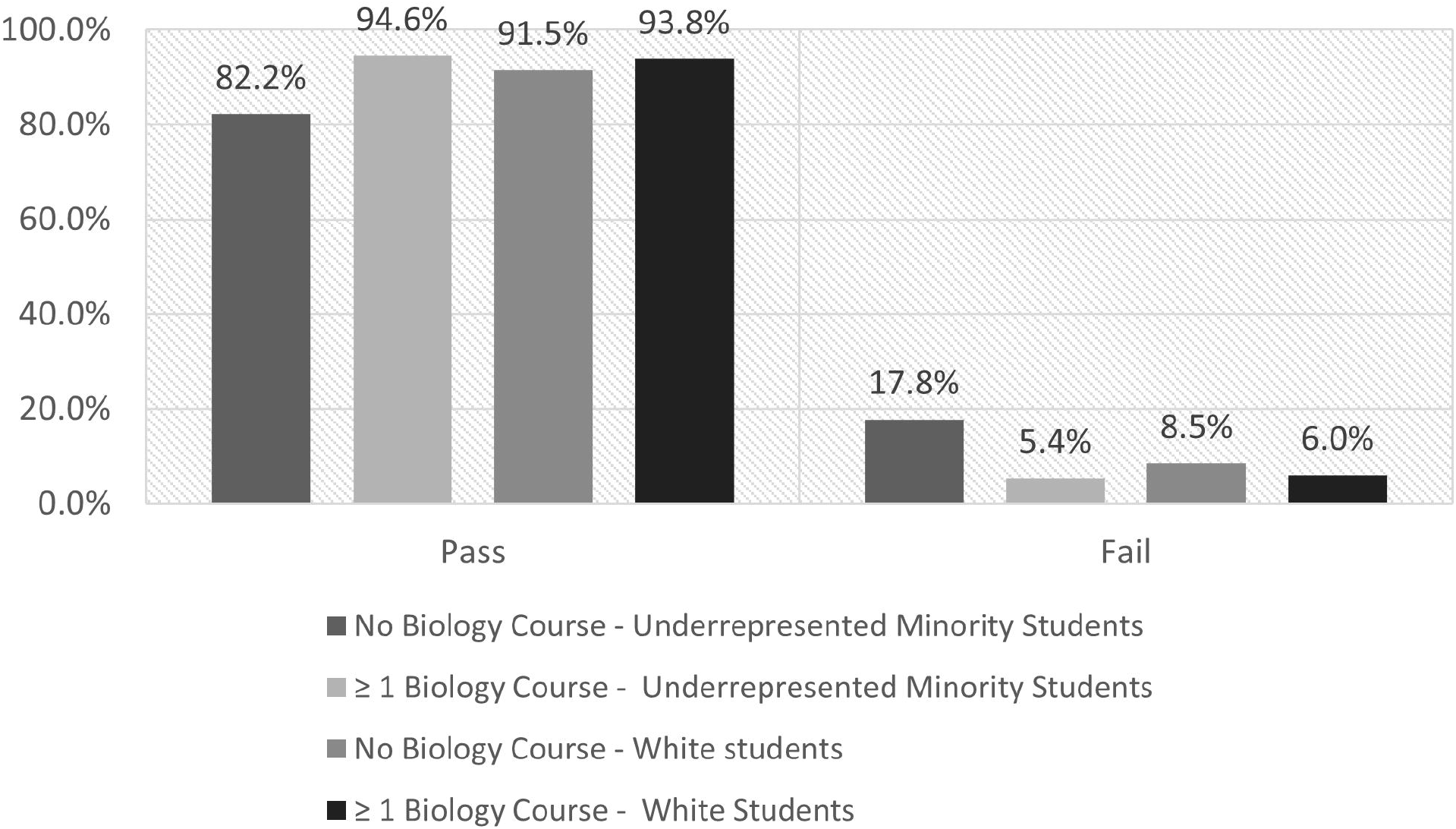

## DISCUSSION

In this study, underrepresented minority students who completed ≥ 1 lab-based college biology course have improved student outcomes in AP as evidenced by an increase in the pass rate from 67% to 78% and a reduction of the failure and withdrawal rate (See Figure 1). While Beeber and Biermann (2007) did not provide statistical analysis, the authors reported the pass rate of at-risk students who took a college biology course before entering AP increased from 66% to 81%. Like the findings presented here, the authors found students taking a biology course had improved course pass rates and reduced failure and withdrawal rates when taking AP (Beeber & Biermann, 2007).

Compared to other studies showing a combined failure and withdrawal rate from 20% to 50% for introductory AP, Madison College’s numbers look pretty good (especially for white students). It would be tempting to be complacent and feel no action is required. However, if we examine these results under the lens of equity, it presents an opportunity to close the achievement gap between white and underrepresented minority students for AP.

How do we close the equity gap at the departmental level? It’s not entirely clear as the solutions aren’t particularly satisfactory or optimal. DeCiccio et al. (2011) provide a framework to understand the challenging choices biology departments face. The first option is the status quo. We cater primarily to high-achieving students or students with privileged backgrounds but fail to fulfill our institutional mission to serve all students – including historically underserved groups (DeCiccio, Kenny, & Lippacher, 2011). The second option is to loosen curriculum and grading standards, but that would create alumni who would not have the tools to succeed in their professional programs and state board exams (DeCiccio, Kenny, & Lippacher, 2011). The last option would be to require foundational courses before enrolling in AP, but this costs more money in additional classes and materials and lengthens the time students must be in the program (DeCiccio, Kenny, & Lippacher, 2011). Despite the difficult choices that biology departments must make, at-risk and underrepresented minority students have innate skills that would make them excellent healthcare professionals, even though some of these students may struggle academically with introductory AP courses. Because we as educators want these students to succeed in their chosen STEM or healthcare field, I concur with DeCiccio et al. (2011) that strategies should include providing remedial coursework or directing students at-risk of failure to comparable programs.

Based on these results, I believe departments should consider requiring an introductory biology course before enrolling in AP. This can be done as straight forward as requiring a lab-based biology course, requiring students to choose between lab-based course or a non-lab-based course, or requiring students to take a placement test before enrollment.

Community colleges are well-positioned to be the vanguard in increasing underrepresented minority representation in STEM and healthcare fields. For example, community colleges train nearly 60% of all new nurses and 80% of new paramedics, respiratory and physical therapists (Kirkwood & Riegelman, 2011; Skillman, Keppel, Paterson, & Doescher, October 2012). There are a few reasons why increasing underrepresented minority representation in these fields would positively impact society and the economy. First, the Association of American Medical Colleges (AAMC) forecasts a shortage of up to 112,00 physicians in the

United States. Similarly, due to high turnover, nurse burnout, and an aging workforce, Haddad et al. project 11 million additional nurses will be needed to avoid further nursing shortages (Haddad, Annamaraju P, & Toney-Butler, 2020). Second, studies have shown that minority patients have improved medical outcomes and report higher trust and increased willingness to get medical procedures when receiving care from African-American medical professionals (Benkert R, 2009; Alsan, 2019). Lastly, medical and health-allied fields typically lead to high-earning careers that can potentially aid in escaping generational poverty and help bridge the wealth gap (Yeung WJ, 2008; Hardaway CR, 2009; Cheng TL, 2016).

## LIMITATIONS

There are a few limitations to this study. First, while the results presented here uncovered a strong association (*p* ≤ 0.008) with prior lab-based college biology coursework and student success in AP for underrepresented minority students, when the withdrawal rates were included, the significance of the results were reduced (*p* = 0.058). Second, like others have noted (Sturges & Maurer, 2013), this study is from a single institution and has limited generalizability. For example, African-American students in the dataset skewed older than a prior study mainly examining non-Hispanic White health science students in Australia (Mills C, Heyworth, Rosenwax, Carr, & Rosenberg, 2009). It is unclear whether this finding on age is generalizable to other African-American populations in a community college setting or a regional or geographical artifact. Lastly, the complete dataset for external GPA and standardized test scores for all students was not available for study nor important factors such as motivation and support, family income, or first-generation status considered. Thus, future directions should include more research that focuses on historically underserved and underrepresented minority students at the community college level as well as high-power statistical studies analyzing the relationship between prior college biology coursework and student success in AP.

## Supporting information

Supplemental Figures of SPSS output

