## Supplemental Figures of SPSS output for "Prior College Biology Coursework Increases Student Outcomes for Underrepresented Minority Students studying Anatomy and Physiology"

Figure S1. Chi-square analysis of pass, failure, and withdrawal rates of non-Hispanic White Students.

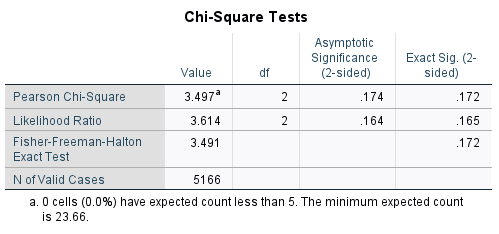

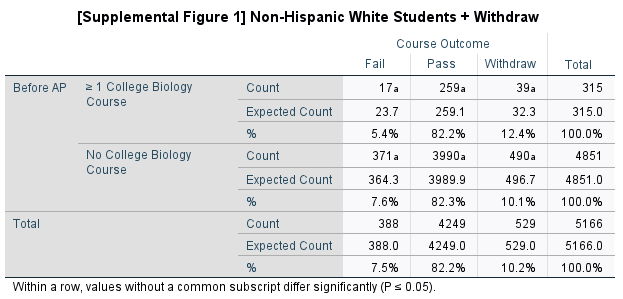

Figure S2. Chi-square analysis of pass, failure, and withdrawal rates of non-Hispanic White Students.

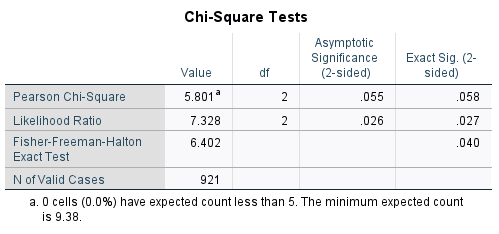

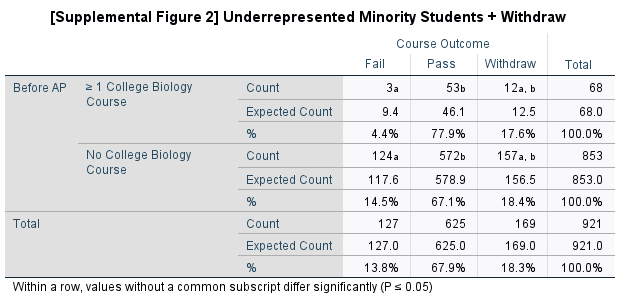

Figure S3. Chi-square analysis of pass and failure rates non-Hispanic White students

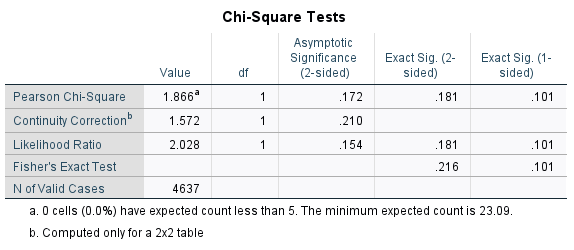

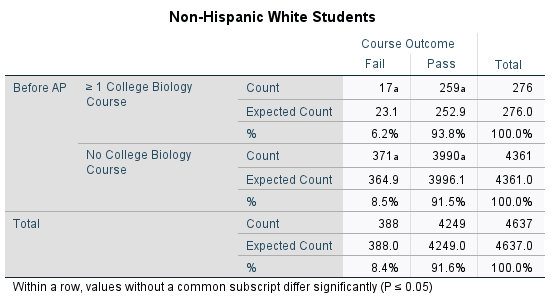

Figure S4. Chi-square analysis of pass and failure rates of Underrepresented Minority students

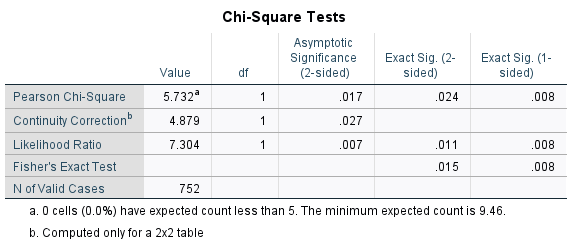

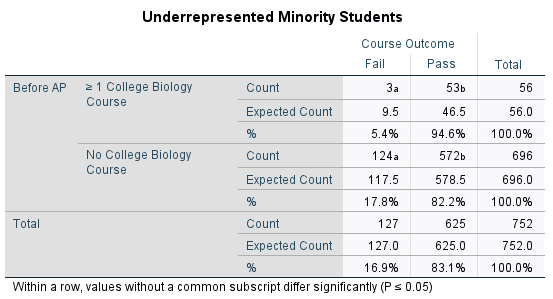
